# Response of *Posidonia oceanica* (L.) Delile and its associated N_2_ fixers to different combinations of temperature and light levels

**DOI:** 10.1101/2021.08.13.456246

**Authors:** Manuela Gertrudis García-Márquez, Víctor Fernández-Juárez, José Carlos Rodríguez-Castañeda, Nona S. R. Agawin

## Abstract

Ocean warming and water turbidity are threats for the persistence of seagrass meadows and their effects on the productivity of seagrasses and the functioning of their associated microorganisms have not been studied extensively. The purpose of this study was to assess the effects of different light levels and temperatures on *Posidonia oceanica*, the endemic seagrass species in the Mediterranean Sea, and their N_2_ fixing community, which contributes importantly to the nitrogen requirements and high productivity of the plants. Aquarium experiments were conducted in winter when the plants are more vulnerable to changes in temperature, subjecting them to short-term exposures to ambient (15.5 °C) and elevated temperatures (ambient+5.5 °C) and at limited (13 μmol photons m^-2^ s^-1^) and saturating light conditions (124 μmol photons m^-2^ s^-1^). Primary production, chlorophyll content, reactive oxygen species production, polyphenols content, the *nifH* gene expression, N_2_ fixation and alkaline phosphatase activities were measured in different plant tissues. Plants incubated at ambient temperature and high light exhibited enhanced total chlorophyll production and significantly higher gross and net primary production, which were approximately two-fold compared to the rest of the treatments. The oxidative stress analyses revealed increased production of reactive oxygen species in young leaves incubated at ambient temperature and saturating light, while the polyphenols content in top leaves was considerably higher under elevated temperatures. In contrast, N_2_ fixation and alkaline phosphatase rates were significantly higher under elevated temperature and low light levels. The presence of the N_2_ fixing phylotypes UCYN-A, -B and -C was detected through genetic analyses, with UCYN-B demonstrating the highest *nifH* gene transcription levels at elevated temperatures. These findings emphasize the significant role of irradiance on the productivity of *P. oceanica* and the temperature dependence of the N_2_ fixation process in winter.

## 1 Introduction

*Posidonia oceanica* (L.) Delile is an endemic and dominant seagrass species in the Mediterranean Sea, where it forms extensive meadows from the surface down to a maximum of about 45 m depth (Procaccini et al., 2003; Boudouresque et al., 2006). Seagrass meadows play major ecological roles by enhancing biodiversity, supporting high productivity, protecting the geomorphology of the coastline, sequestering global oceanic carbon and providing a buffering effect against ocean acidification (Duarte et al., 2005; Barbier et al., 2011; Fourqurean et al., 2017; Chou et al., 2018). However, seagrass ecosystems are currently suffering from a worldwide regression in response to several environmental stressors (Orth et al., 2006; Waycott et al., 2009; Marbà and Duarte, 2010). For instance, the reduction in the surface coverage of *P. oceanica* has been reported to be 34% in the past 50 years (Telesca et al., 2015). Considering that *P. oceanica* is a climax, slow-growing seagrass species, natural and anthropogenic perturbations can be particularly critical, as their recovery from perturbations can be very slow or may not recover at all (Serrano et al., 2011). Among the threats that these ecosystems are facing, eutrophication from waste waters and aquaculture, shoreline constructions, anchoring and trawling, dredging, introduced species, and climate change (warming and sea-level rise) are considered major causes of the decline of *P. oceanica* meadows over the last decades (Boudouresque et al., 2009; Champenois and Borges, 2018). Most of these impacts potentially or ultimately reduce water transparency and, therefore, the quality and quantity of the irradiance reaching the seagrass canopy (Duarte et al., 2004; Orth et al., 2006). Epiphytic and planktonic algal accumulations from excess anthropogenic nutrients, increased sediment run-off, and resuspension of bottom sediments are primary causes of reduced underwater irradiance in coastal areas (Orth and Moore, 1983; Cambridge et al., 1986; Onuf, 1994). Ocean warming is also regarded as one of the most severe factors of global climate change, expected to cause, under extreme greenhouse gas emission scenarios, the rise of ocean surface temperatures between 2.6 °C and 4.8 °C by 2100, along with an increased amplitude and duration of heat waves (abnormally warm seawater episodes) (IPCC, 2014). These changes are predicted to have serious repercussions in the Mediterranean Sea given its confined nature, which makes it more susceptible to temperature increases, with warming occurring at significantly higher rates compared to open oceans (Diffenbaugh et al., 2007; Vargas-Yáñez et al., 2008; Calvo et al., 2011).

Generally, light and nutrients comprise the source of energy and matter needed for the growth of seagrasses, while temperature regulates biochemical processes involved in photosynthesis and respiration, thus, predominantly controlling the annual and seasonal production patterns of seagrasses (Lee and Dunton, 1996; Zupo et al., 1997; Lee et al., 2005, 2007). Water temperature and irradiance are usually correlated and display similar seasonal trends, making it difficult to isolate both environmental parameters in relation to seagrass growth and production (Kaldy and Dunton, 2000; Kaldy, 2006). Light requirements for seagrasses are unusually high, being approximately 10-37% of surface irradiance, compared to the 0.1-1% needed for most of the other marine macrophytes, that is partially attributed to inefficient carbon concentrating mechanisms for photosynthesis (Invers et al., 2001; Larkum et al., 2006; Zimmerman, 2006). In view of this, seagrasses are highly vulnerable to the deterioration of water clarity, which is evidenced through the reported large-scale losses of meadows worldwide (Dennison et al., 1993; Onuf, 1994; Short and Wyllie-Echeverria, 1996; Erftemeijer et al., 2006). Hence, understanding the light thresholds for seagrass survival is fundamental for an effective management of these valuable habitats (York et al., 2013). Elevated temperatures entail a grand risk of local extinction for cold-adapted plants, such as Mediterranean seagrasses, as they have visibly manifested physiological symptoms of heat stress and reduced fitness (Beca-Carretero et al., 2018; Marín-Guirao et al., 2018). The consequences of heat waves can be notably damaging on seagrass meadows, promoting shoot mortality and population decline when critical temperature thresholds are surpassed (Díaz-Almela et al., 2009; Marbà and Duarte, 2010; Jordà et al., 2012). In addition, thermal stress can induce the acceleration of the respiration over photosynthesis rates (Collier and Waycott, 2014; Marín-Guirao et al., 2018), and affect important life history events, like reproduction, through increased flowering intensity (Ruiz et al., 2018; Marín-Guirao et al., 2019).

Several studies have been carried out in order to determine the effects of light reduction on *P. oceanica*, comparing its response along bathymetric/spatial gradients (Alcoverro et al., 2001; Ruiz and Romero, 2003; Dattolo et al., 2014) and experimentally though the modification of the light environment with shading screens (Ruiz and Romero, 2001; Mazzuca et al., 2009; Serrano et al., 2011; Gacia et al., 2012). Investigations on the impacts of sea warming, considering climate change scenarios, have focused on its isolated effects (Marbà and Duarte, 2010; Guerrero-Meseguer et al., 2017; Hernán et al., 2017; Ruiz et al., 2018; Traboni et al., 2018) and the interaction with other stressors (Hendriks et al., 2017; Ontoria et al., 2019; Agawin et al., 2021). However, research on the combined effects of ocean warming and water turbidity in *P. oceanica* meadows remains relatively scarce and understanding them is crucial for the future management of these coastal ecosystems. Moreover, no studies have been done so far on the effect of these factors on the microorganisms associated with *P. oceanica*, such as the nitrogen (N_2_) fixers.

Biological N_2_ fixation, defined as the enzymatic reduction of atmospheric N_2_ to ammonium equivalents, takes part in significantly sustaining the high productivity of *P. oceanica* in the oligotrophic Mediterranean Sea (Garcias-Bonet et al., 2019). Seagrasses harbor diverse communities of epi- and endophytic bacteria associated with their leaves, roots and rhizomes, that can enhance plant growth through increased nutrient availability, for instance, via N_2_ fixation or by mineralizing organic compounds (Uku et al., 2007; Cole and McGlathery, 2012; Garcias-Bonet et al., 2016). In particular, it has been reported that N_2_ fixation processes associated with the phyllosphere of *P. oceanica* could potentially supply the total nitrogen demand of the plant (Agawin et al. 2016; 2017). The enzyme complex that catalyzes the biological N_2_ fixation process is the nitrogenase, which is composed of two proteins: conventionally, the iron protein or nitrogenase reductase, and the molybdenum iron protein or nitrogenase (Hamisi, 2010). The *nifH* gene encodes the iron protein and the *nifDK* genes encode the molybdenum iron protein (Rubio and Ludden, 2002). Sequencing the *nif* genes, with *nifH* being the most sequenced and marker gene of choice, has allowed studying the phylogeny, diversity, and abundance of N_2_-fixing microorganisms (Gaby and Buckley, 2012). A significant diversity has been identified among marine diazotrophs, with filamentous organisms including primarily *Trichodesmium* sp. and the diatom symbiont *Richelia intracellularis*, and highly diverse unicellular diazotrophs that comprise Cyanobacteria, Proteobacteria, and Archaea (Moal et al., 2011). At present, three groups of unicellular N_2_-fixing cyanobacteria (UCYN) have been described: UCYN-A, B, and C (Zehr et al., 2001; Foster et al., 2007). Groups B and C are nanoplanktonic cells (2-10 μm) closely related to the cultivated strains *Crocosphaera watsonii* and *Cyanothece* sp., respectively (Church et al., 2005; Foster et al., 2007); whereas members of Group A are of picoplanktonic size (0.7-1.5 μm) and uncultivated up until now (Biegala and Raimbault, 2008; Goebel et al., 2008).

Nitrogenase activity is influenced by a combination of environmental factors that differ depending on the geographic region and diazotroph community composition (Mahaffey et al., 2005). Several studies have suggested a strong temperature dependence of the N_2_ fixation process at the enzymatic level, and a positive correlation to irradiance (Welsh, 2000; Brauer et al., 2013; Agawin et al., 2017; Garcias-Bonet et al., 2019). Furthermore, as the nitrogenase proteins require iron (Fe), the availability of both phosphorus (P) and Fe are factors that could limit or co-limit the N_2_ fixation process in some areas of the oceans (Sañudo-Wilhelmy et al., 2001; Karl et al., 2002; Mills et al., 2004). Aquatic primary producers usually contain external alkaline phosphates; enzymes capable of hydrolyzing organic phosphorus compounds (monoester phosphates), which liberates inorganic phosphorus and increases the availability of this nutrient for growth (Kuenzler and Perras, 1965; Martínez-Crego et al., 2006). Thus, the measurement of alkaline phosphatase activity has been employed as an indicator of phosphorus limitation and deficiency in algae and seagrasses (Pérez and Romero, 1993; Invers et al., 1995; Steinhart et al., 2002; Fernández-Juárez et al., 2019).

The purpose of this study was to assess the response *P. oceanica* and its N_2_ fixing community to different combinations of temperature and light levels, in terms of primary production and respiration rates, chlorophyll content, alkaline phosphatase activity, oxidative stress indicators and N_2_ fixation activities of the diazotrophs associated with different plant tissues. The experiment was performed during winter, when the plants are thermally more vulnerable to temperature increases (Agawin et al., 2021).

## 2 Materials and Methods

### 2.1 Sampling and experimental design

To assess the effects of warming and deteriorating light conditions on *Posidonia oceanica* and their N_2_ fixing community, aquarium experiments were conducted in winter simulating combinations of present and future temperatures (IPCC, 2007) with two light conditions. Limited (13 μmol photons m^-2^ s^-1^) and saturating (124 μmol photons m^-2^ s^-1^) light levels, based on the photosynthesis-irradiance parameters documented in the literature for shallow *P. oceanica* meadows during winter (Alcoverro et al., 1998; Lee et al., 2007), were combined factorially with the ambient temperature corresponding to the time of the collection (15.5 °C) and 5.5 °C warmer (21 °C). The plants were carefully collected from the coast of Llucmajor (2°44’22.65”E, 39°27’2.36”N; Majorca, Spain; Fig. 1) in December 2020, through SCUBA diving at a depth between 4 to 6 m. Seawater was also collected and immediately prefiltered through a 10 μm nylon Nitex filter, of which 8 L were added to each of 12 aquaria with 9 L of capacity. Three replicate aquaria were employed per treatment and 8-10 shoots of *P. oceanica*, with roots and part of the rhizome attached, were placed in each aquarium without sediments. The cut end of the horizontal rhizome of each plant was sealed using a non-toxic underwater D-D AquaScape epoxy to maintain gas pressure inside the rhizome. The experiment was performed in a temperature-controlled room, with a duration of 18 days, in between which the seawater was replaced to avoid nutrient limitation in the aquaria. The temperature treatments were achieved by respectively connecting each aquarium to water chillers (HAILEA HC-130A) with a continuous circuit of water and heaters (Aquael EasyHeater 25 W), with the desired temperature previously configured in the devices. Aquaria were illuminated by diode lamps (Aquael Leddy Slim Sunny 5 W) installed above, set to 11:13 h light: dark cycles and delivering incident PAR light levels at the seagrass canopy according to the treatments assigned. The partial pressure of carbon dioxide (CO_2_) was adjusted through bubbling with an air-CO_2_ mixture. Atmospheric air was first scrubbed by soda lime to remove all CO_2_ and then mixed with pure CO_2_ from a bottle using mass flow controllers (Aalborg). To achieve present-day pCO_2_ levels, gases were mixed to 435 ppm pCO_2_ in mixing bottles filled with marbles to assure the homogenization of gases. In each aquarium, the resulting mixture was regulated by a flow meter with a volume of 2.5 L min^-1^, and a flux diffuser was placed at the extremes of each tube to release the gases in diffused form.

**Figure 1.**
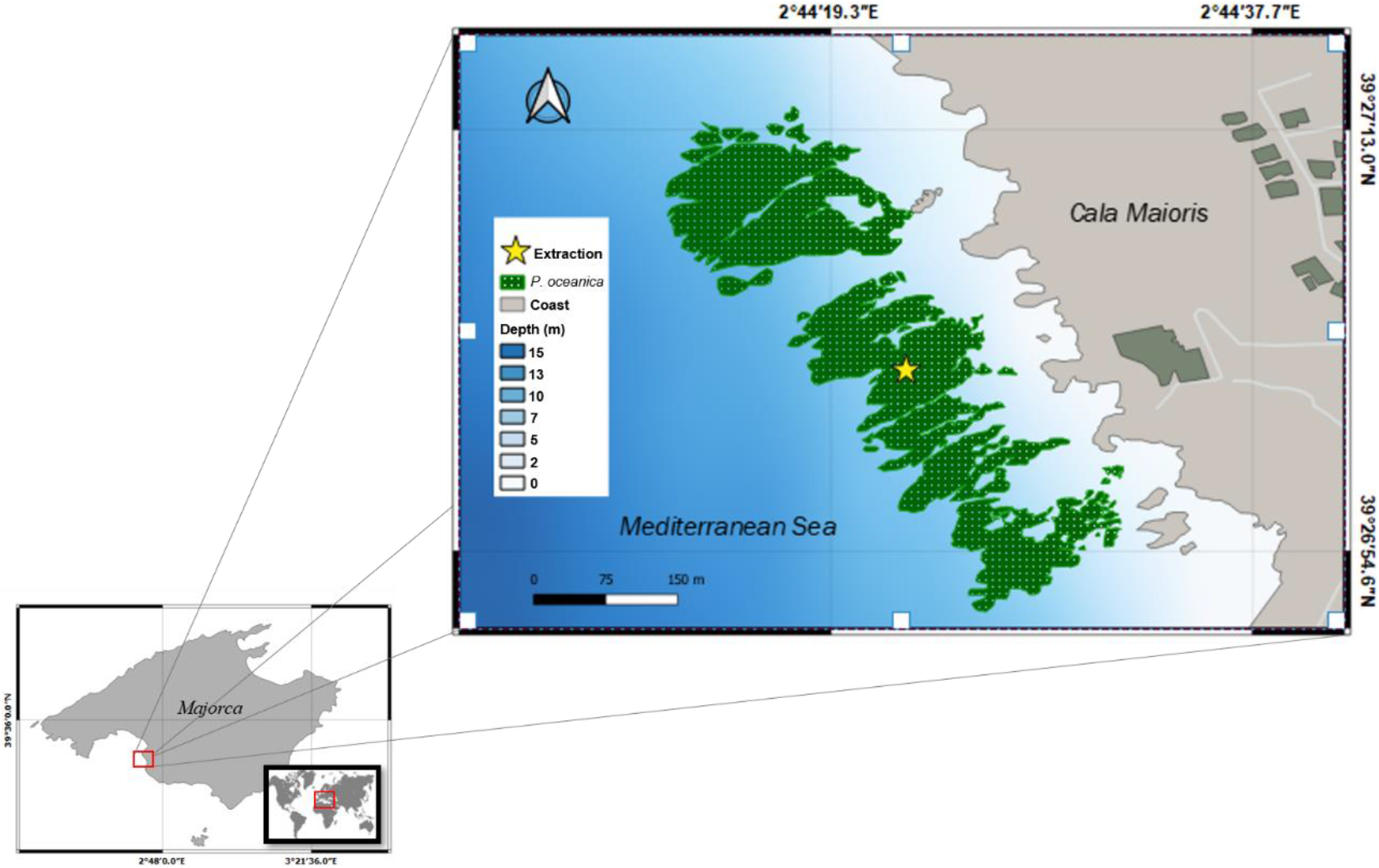
Geographical location of the collection site of *Posidonia oceanica* in the coast of Llucmajor, Majorca, Spain.

### 2.2 Physicochemical parameters

Temperature (IKS-Aquastar) and pH (ENV-40-pH, calibrated with 4.0 and 7.0 pH NBS standards) were continuously monitored and recorded at 30 min intervals using sensors, connected to a D130 data logger (Consort) and computer. The daily average photosynthetically active radiation (PAR) was monitored with light loggers (HOBO), which were positioned in the surface water of the aquaria. Due to a limited number of sensors and light loggers, only two replicates per treatment could be measured simultaneously for the indicated parameters. Before, in the middle and after the incubations, water samples from each aquarium were taken for the determination of nitrite (NO_2_^-^), nitrate (NO_3_^-^), ammonia (NH_4_^+^), phosphate (PO_4_^3-^) and total dissolved phosphorus (TDP) concentrations. The samples were filtered through sterile polypropylene filter holders (0.2 μm) using a peristaltic pump (Geotech Geopump) and kept frozen until analyzed. The inorganic nutrients samples (NO_2_^-^, NO_3_^-^, NH_4_^+^, PO_4_^3-^) were stored in polypropylene tubes, while samples for TDP were deposited in borosilicate Scott bottles. NO_2_^-^ concentrations were quantified following the spectrophotometric method of Strickland and Parsons (1972), and a modified protocol based on Knap et al. (1997) and Weber-Shirk et al. (2001) was applied for PO_4_^3-^ determination. TDP concentrations were also analyzed using the latter method after persulfate digestion (Bronk et al., 2000). NO_3_^-^ content was determined by flow injection analysis as described by Diamond (2003) and NH_4_^+^ was measured according to the modified fluorometric method of Horstkotte and Duarte (2012).

### 2.3 Estimation of primary production and respiration rates

Dissolved oxygen (DO) concentrations were determined spectrophotometrically by the modified Winkler method, according to the protocol described by Labasque et al. (2004). For each aquarium, four Exetainer vials (initial values, *n*=4) (12 mL) and two light and dark 125 ml Winkler bottles were filled with water from the aquarium filtered through sterile polypropylene filter holders (0.2 μm) using the peristaltic pump, taking care to avoid bubbles or turbulence when filling. The second youngest leaf of each of four independent shoots per aquarium was selected, cut into a 5 cm segment from the top and, if necessary, epiphytes were scraped off. Each leaf segment was inserted into the light and dark Winkler bottles to incubate for 3 hours inside their respective aquariums. For phyllosphere measurements, two Erlenmeyer flasks were filled with 480 ml of filtered water from each aquarium and autoclaved, then a *P. oceanica* shoot without roots and rhizomes was introduced per flask and incubated as previously mentioned. After the incubation period, Exetainers were filled with the water from the Winkler bottles and flasks until they overflowed, using the syringe with the attached tube to avoid gas exchange as much as possible. Immediately, 80 μL of MnCl_2_ (3 M) and 80 μL of NaOH (8 M) and NaI (4 M) were added in the vials. Exetainers were tightly closed, agitated, and kept in cold and dark conditions until DO determination (between 24 and 48 h). DO concentrations were estimated spectrophotometrically at 466 nm after adding 80 μL H_2_SO_4_ (10 M). The increase or decrease of DO concentrations during the incubation period provided measures of net primary production (NPP) and respiration (R) in the light and dark bottles, respectively. Then, the gross primary production (GPP) was calculated by summing the net photosynthetic rates obtained with the rate of dark respiration (*GPP* = *NPP* + *R*). The estimated changes in DO from the Erlenmeyer flasks provide the NPP of the *P. oceanica* phyllosphere. These values were normalized to incubation time, volume of water and the dry weight of the incubated tissue (μmol O_2_ g DW^-1^ h^-1^).

### 2.4 Determination of chlorophyll concentrations

Leaf chlorophyll concentrations in duplicate *P. oceanica* shoots from each aquarium were measured following Agawin et al. (1996). Extraction of chlorophyll a and b from the seagrass leaves was done by grinding about 0.1 to 0.3 g wet weight of the second youngest leaf per shoot, with a mortar and pestle in 96% ethanol. After extraction in the dark for 12 h, the suspensions were centrifuged at 2800×g for 10 min. Absorbances were measured at 665 and 649 nm using a Cary-50 Conc-UV Visible spectrophotometer. Afterwards, chlorophyll a and b concentrations were determined using the formula of Wintermans and De Mots (1965).

### 2.5 Quantification of alkaline phosphatase activity

Alkaline phosphatase activity (APA) was evaluated through a fluorometric assay, in which the hydrolysis of the fluorogenic substrate (S) 4-methylumbelliferyl phosphate (MUF-P, Sigma-Aldrich) to 4-methylumbelliferyl (MUF) was measured (Fernández-Juárez et al., 2019). The second oldest and youngest leaf of each of two independent shoots per aquarium were selected and cut into a 5 cm segment from the top. From each of the two independent shoots, 5 cm piece of unrinsed rhizomes and roots were also extracted. The leaf and root segments were inserted into 15 ml centrifuge tubes with 10 ml of filtered and autoclaved water from their respective aquariums, while the rhizomes were introduced into 50 ml Falcon centrifuge tubes with 40 ml of the water. Then, the MUF-P reagent at 2 μM of final concentration was added to each tube. After 1 h incubation in darkness at room temperature, APA was measured in a microtiter plate that contained borate buffer at pH 10 (3:1 of sample:buffer). The MUF production (fmole MUF cell-1 h-1) was measured with a Cary Eclipse spectrofluorometer (FL0902M009, Agilent Technologies) at 359 nm (excitation) and 449 nm (emission), and using a calibration standard curve with commercial MUF (Sigma-Aldrich).

### 2.6 Reactive oxygen species production

Prior to biochemical analysis, *P. oceanica* leaves were carefully separated from the epiphytes. The leaf segments per aquarium were washed with distilled water to eliminate salt residues and triturated in a mortar with pestle in the presence of liquid nitrogen. The samples were homogenized in five volumes (w/v) of 50 mM Tris-HCl buffer, 1 mM EDTA, pH 7.5. Then, the solutions were homogenized in ice employing a homogenizer, with a velocity set between 4-6, for a few minutes. Homogenates were centrifuged at 9000×g at 4 °C for 4 min to remove cell debris, nuclei and mitochondria and the supernatants were used for biochemical assays. All biochemical analyses were expressed per mg protein, measured by using the colorimetric Thermo Scientific Coomassie (Bradford) Protein Assay Kit with Bovine Serum Albumin (BSA) as a standard. The reactive oxygen species (ROS) production was measured using the molecular probe 2’,7’-dichlorofluorescein diacetate (DCFH-DA; Sigma) in culture media (ASN-III+C Turks Island salts 4× or BG11_0_), which was added to a 96-well microplate (Thermo Scientific) containing the supernatant samples (final concentration of probe at 15 μg ml^-1^). This compound is intracellularly hydrolyzed by esterases to non-fluorescent 2’,7’-dichlorodihydrofluorescin (DCFH), which is subsequently oxidized by ROS to highly green fluorescent 2’,7’-dichlorodihydrofluorescein (DCF) (Kumar et al., 2018). The fluorescence was measured at 25 °C in a FLx800 Microplate Fluorescence Reader (BioTek Instruments, Inc.) for 1 h, with an excitation of 480 nm and emission of 530 nm. The measurements were obtained from the slope of the linear regression between the fluorescence readings and time, and expressed as arbitrary units (AU). DCFH-DA was added in ASN-III+C Turks Island salts 4× or BG11_0_ without sample as blanks under the same conditions stated above.

### 2.7 Phenolic compounds quantification

The total phenolic content of the *P. oceanica* extracts was estimated by the Folin-Ciocalteau colorimetric assay (Singleton et al., 1999). Briefly, 10 μl of the extract sample was mixed with 10 μl of 2 N Folin-Ciocalteu reagent, 50 μl of 20% (w/v) sodium carbonate (Na_2_CO_3_) and 250 μl of distilled water. After incubation at room temperature for 90 min, absorbance was measured at 760 nm (UV-visible spectrophotometer Cary 100 Conc, Varian). A calibration curve was built by using tyrosine as the standard and the total phenolic content was expressed as mg of tyrosine/mg of protein. All determinations were carried out in duplicate per aquarium.

### 2.8 Measurement of N_2_ fixation rates

N_2_ fixation rates were measured in the different plants tissues of *P. oceanica* using the acetylene reduction assay (ARA) (Stal, 1988; Capone, 1993; Agawin et al., 2014). The second oldest leaf, rhizomes and roots of each of two independent shoots per aquarium was selected and cut into a 5 cm segments. Additional 5 cm pieces of roots were also extracted from independent shoots for surface-sterilization by a series of sterilization steps (i.e. 99% ethanol 1 min; 3.125% NaOCl 6 min; 99% ethanol 30 s; autoclaved GF/F filtered seawater final washing; Coombs and Franco 2003), in order to measure root endophyte N_2_ fixation rates. Each plant tissue was inserted into its respective incubation vial. Leaves and roots were inserted into 10 ml gas chromatograph (GC) vials and the rhizomes into 50 ml Falcon centrifuge tubes. Each incubation vial or tube was humidified with 1 ml (for the GC vials) and 2.5 ml (for the Falcon tubes) sterilized GF/F filtered seawater. All vials and tubes were capped with gas-tight septum ports. Vials and tubes containing the rhizomes and roots were flushed with helium gas for 1 min to obtain anoxic conditions. Each incubation vial or tube was injected with volume of acetylene gas at 20% (v/v) using gas-tight Hamilton syringes, and then incubated for 3 h in their respective aquarium. Immediately after the incubation time, 10 ml of headspace was taken using a gas-tight Hamilton syringe from the incubation vials or tubes, transferred to Hungate tubes and sealed with hot melt glue (SALKI, ref. 0430308) to avoid possible gas losses as much as (Agawin et al., 2014). Ethylene and acetylene were determined using a gas chromatograph (7890A, Agilent Technologies) equipped with a flame ionization detector. The column was a Varian wide-bore column (ref. CP7584) packed with CP-PoraPLOT U (27.5 m length, 0.53 mm inside diameter, 0.70 mm outside diameter, 20 μm film thickness). Helium was used as carrier gas at a flow rate of 30 ml min^-1^. Hydrogen and airflow rates were set at 30 ml min^-1^ and 365 ml min^-1^, respectively. The split flow was used so that the carrier gas flow through the column was 4 ml min^-1^ at a pressure of 5 psi. Oven, injection and detector temperatures were set at 52°C, 120°C and 170°C, respectively. The amount of ethylene produced was obtained following the equations in Stal (1988). The acetylene reduction rates were converted to N_2_ fixation rates using a factor of 4:1 (C_2_H_4_:N_2_ reduced; Jensen and Cox 1983) and reported per g dry weight of plant biomass incubated. The dry weight of the plant parts was determined by drying the plant parts at 60°C for 24 h (Short and Duarte, 2001).

### 2.9 Quantification of the *nifH* gene expression in the phyllosphere of *Posidonia oceanica*

After the incubations, for the extraction of the epiphytic community in *P. oceanica*, the leaf segments from each aquarium were placed onto clean glass slides and scraped on both sides with new sterile disposable scalpel blades (#10). The epiphytes obtained per aquarium were transferred into eppendorf tubes with 1 ml of phosphate buffered saline (PBS) solution, in order to remove salt residues that could interfere during the RNA extraction process, and then centrifuged at 13000×g for 15 min. RNA extraction and purification was done with the Plant/Fungi Total RNA Purification Kit (Norgen, Cat. 25800, 31350, 25850), following the manufactures protocol. The quality and quantity of the extractions (absence of DNA and protein contaminations) were assessed using NanoDrop (Thermo Fisher Scientific). The expression of the *nifH* gene was assessed by a Reverse Transcription-quantitative Polymerase Chain Reaction (RT-qPCR) as described by Goebel et al. (2010), Moisander et al. (2010) and Turk-Kubo et al. (2012), considering primer sets designed for N_2_ fixing communities belonging to the Groups A, B and C of unicellular cyanobacteria, the filamentous cyanobacteria genera *Trichodesmium*, and alpha-proteobacteria. The assays were performed in the LightCycler 480 Instrument II - Roche Life Science, using the Luna Universal One-Step RT-qPCR Kit. All RT-qPCR reactions were carried out in triplicate to capture intra-assay variability, and each assay included three no-template negative controls for each primer pair. The cycle threshold (CT) values were used to calculate the number of gene copies per sample, based on the standard curves for each primer set, and normalized to the total RNA content.

### 2.10 Data and statistical analyses

Data is presented as mean ± standard deviation of the replicates from the treatments (*n*=3). Prior to the statistical analyses, data were tested for normality using the Shapiro-Wilk (*n*<50) and Kolmogorov-Smirnov (*n*>50) goodness of fit tests, while the homoscedasticity was assessed with Levene’s test, and then log-transformed if necessary. One-way analysis of variance (ANOVA) was used to test the hypothesis that GPP, NPP and respiration rates of *P. oceanica* vary among the different treatments. The effect of the treatments on chlorophyll content was examined through linear mixed models (LMM), including the aquaria as random factor. For the remaining biological parameters (APA rates, ROS production, polyphenols content, N_2_ fixation, and *nifH* expression), LMM were also executed in order to evaluate possible differences among treatments and plant tissues, with the aquaria as random factor, and considering the interaction between fixed factors. Finally, post-hoc analyses were performed with the Tukey test for multiple comparisons of means. The statistical analyses were performed using the R package, version 4.0.3.

## 3 Results

### 3.1 Physicochemical parameters

The mean temperature of the aquaria at ambient and elevated temperature corresponded to 15.70±0.47 and 21.48±0.57 °C, respectively. On average, the low and high light treatments differed, although not significantly, at ambient temperature with 0.86±0.07 °C, and 0.82±0.17 °C under elevated temperature, with the high light treatments reaching slightly higher values in both cases (Fig.2A). The pH of tanks receiving low light exhibited a lower mean (7.82±0.08) compared to those subjected to high light conditions (8.04±0.11). The temporal fluctuations of the pH in all treatments is showed in Fig. 2B, with diurnal changes of approximately 0.09 to 0.42 units. Regarding PAR values, low and high light treatments were daily exposed to an average of 11.69±2.45 and 126.29±8.77 μmol photons m^-2^ s^-1^, respectively (Fig. 2C). In the nutrient analyses performed, a decrease in the NO_3_^-^ and PO_4_^3-^ concentrations of all treatments was evidenced towards the end of the experiment (Table 1), while NO_2_^-^, NH4^+^ and TDP values were lower compared to the initial phase only before the water replacement. Furthermore, at the final stage of the incubations, NO_2_^-^ concentrations were higher in all treatments and the NH4^+^ was higher in aquaria under saturating light conditions, in comparison to the values obtained at the intermediate water replacement. A slight decrease in the PO_4_^3-^ concentrations was observed before the water replacement in the ambient temperature with high light treatment only. On the other hand, the TDP concentrations increased towards the end of the experiment in tanks under ambient temperature with low light and elevated temperature with high light.

**Figure 2.**
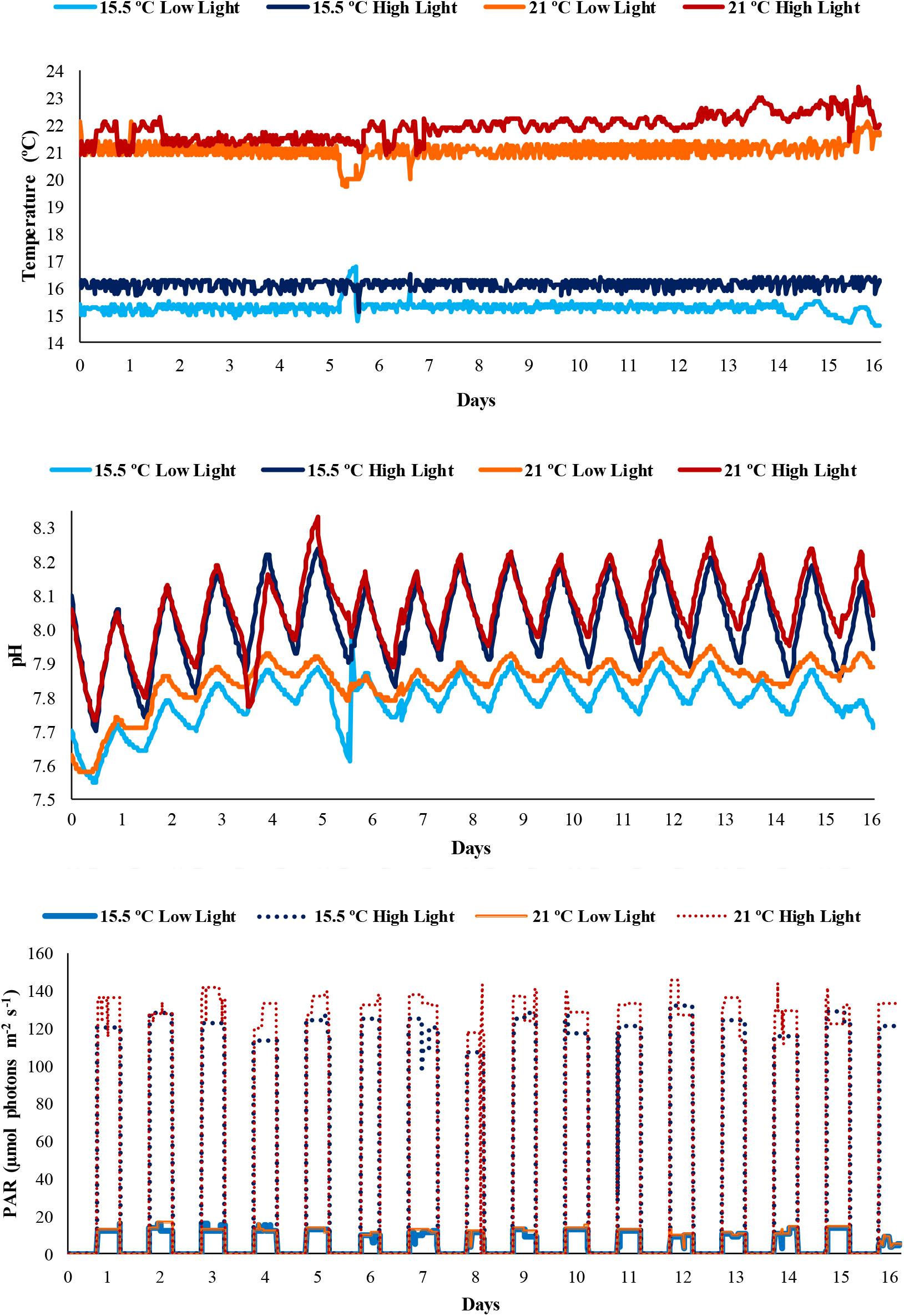
The diurnal variations of **(A)** temperature, **(B)** pH and **(C)** photosynthetically active radiation (PAR) values measured during the experiment.

**Table 1.**
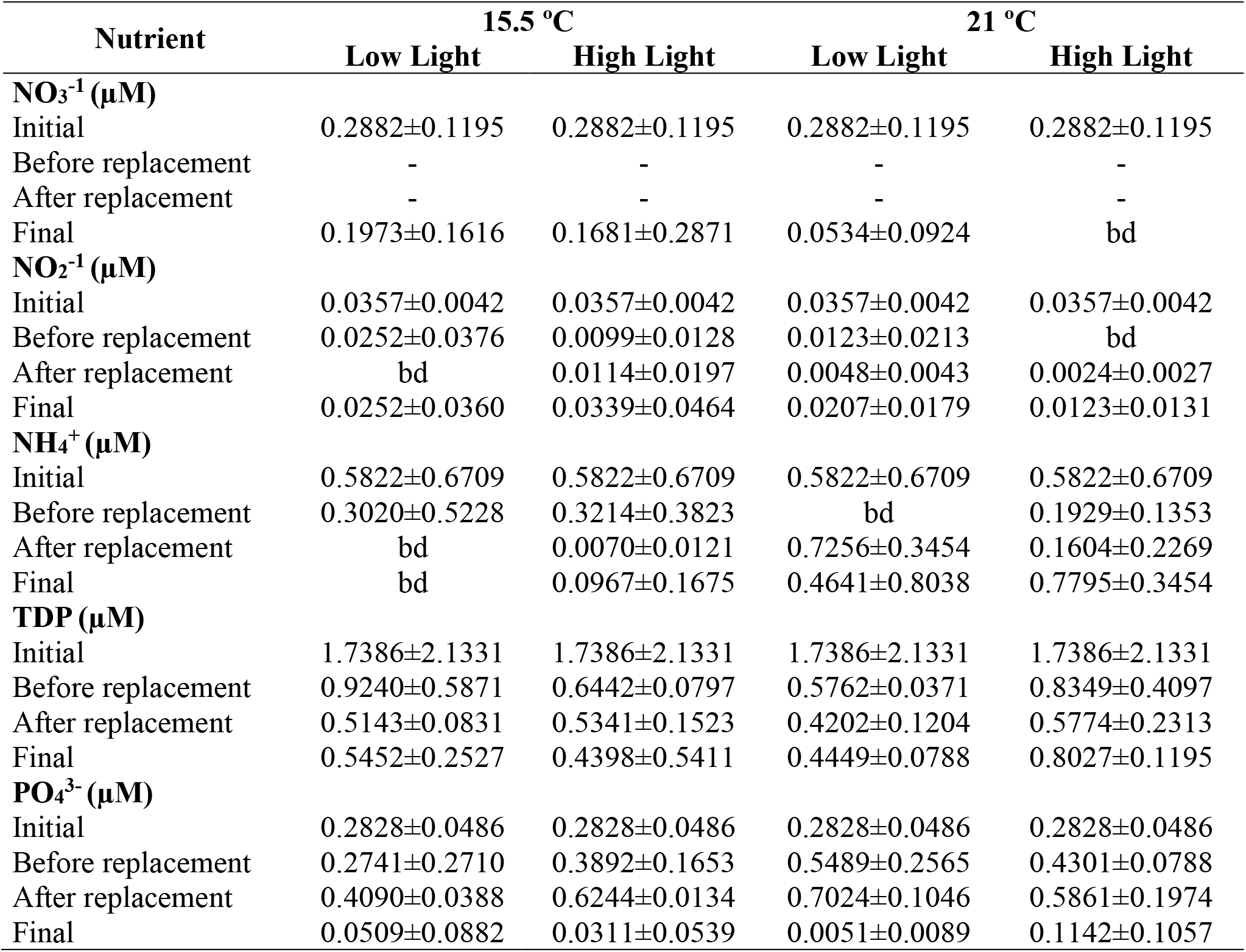
The average (± SD, *n*=3) concentrations of nitrate (NO_3_^-^), nitrite (NO_2_^-^), ammonium (NH_4_^+^), total dissolved phosphorus (TDP) and phosphate (PO_4_^3-^) during the course of the experiment, including the initial at day zero, before and after water replacement at day eight, and final at day 18 (bd: below detection).

### 3.2 Primary production and respiration rates

The average GPP rates of cut leaf segments were significantly higher (*p*<0.05; Table S1, Supplementary Material) under high light conditions at ambient temperature (9.61±1.62 mg O_2_ g DW^-1^ h^-1^) compared with the low light treatments (15.5 °C=5.32±1.84 mg O_2_ g DW^-1^ h^-1^; 21 °C=5.77±0.59 mg O_2_ g DW^-1^ h^-1^) (Fig. 3A). Similar results were obtained for the whole phyllosphere, where the mean NPP rates were significantly higher (*p*<0.01) at saturating light conditions (21 °C=0.90±0.60 mg O_2_ g DW^-1^ h^-1^; 15.5 °C=0.50±0.23 mg O_2_ g DW^-1^ h^-1^) compared to the limited light treatment at elevated temperature (0.11±0.09 mg O_2_ g DW^-1^ h^-1^) (Fig. 3B). The leaves incubated at ambient temperature and high light reached the highest NPP rates (*p*<0.05), with a mean value of 6.07±1.42 mg O_2_ g DW^-1^ h^-1^, which is approximately two-fold in comparison to the remaining treatments. Although the high light treatments exhibited the highest respiration rates, at elevated (−3.64±2.13 mg O_2_ g DW^-1^ h^-1^) and ambient temperatures (−3.54±3.03 mg O_2_ g DW^-1^ h^-1^), respectively, these values did not differ significantly from the remaining treatments (*p*>0.05).

**Figure 3.**
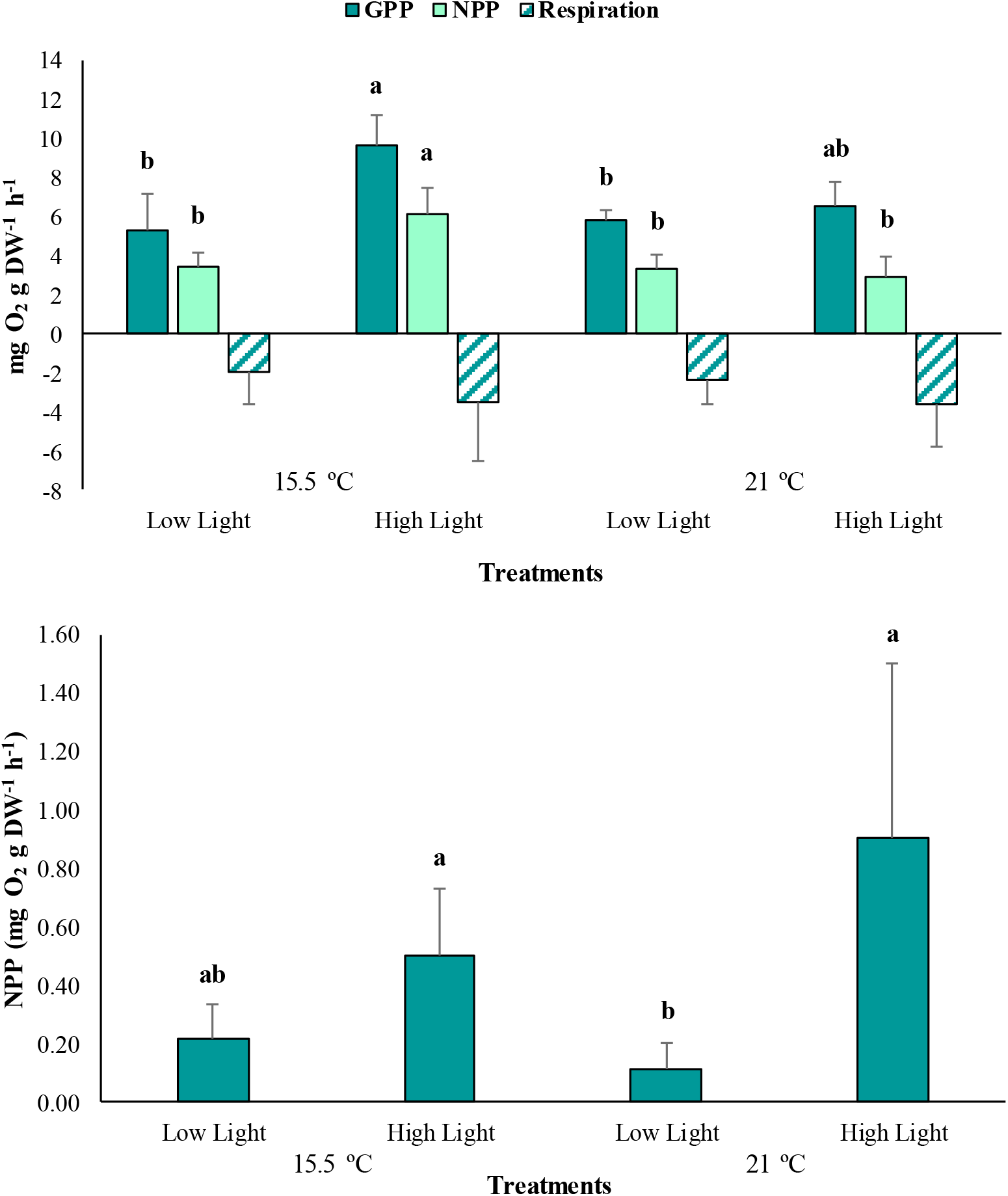
**(A)** The average gross primary production (GPP), net primary production (NPP) and respiration rates of *Posidonia oceanica* cut leaf segments in the different treatments (*n*=3). **(B)** The average NPP rates of the *P. oceanica* phyllosphere in the different treatments (*n*=3). The error bars represent ± SD. Different letters denote significant differences (*p*<0.05) among treatments (Tukey’s post-hoc test following the respective analysis of variance/deviance in Table S1, Supplementary Material).

### 3.3 Chlorophyll concentrations

Total chlorophyll concentrations, as well as chlorophyll *a* and *b*, demonstrated a corresponding trend with primary production, with considerably enhanced mean values (*p*<0.001; Table S2, Supplementary Material) under high light conditions, at ambient (Total Chl=329,67±47.91; Chl *a*=194.46±22.91; Chl *b*=135.21±27.83 μg g WW^-1^) and elevated (Total Chl=259.98±26.49; Chl *a*=152.88±13.98; Chl *b*=107.10±17.23 μg g WW^-1^) temperatures, respectively (Fig. 4).

**Figure 4.**
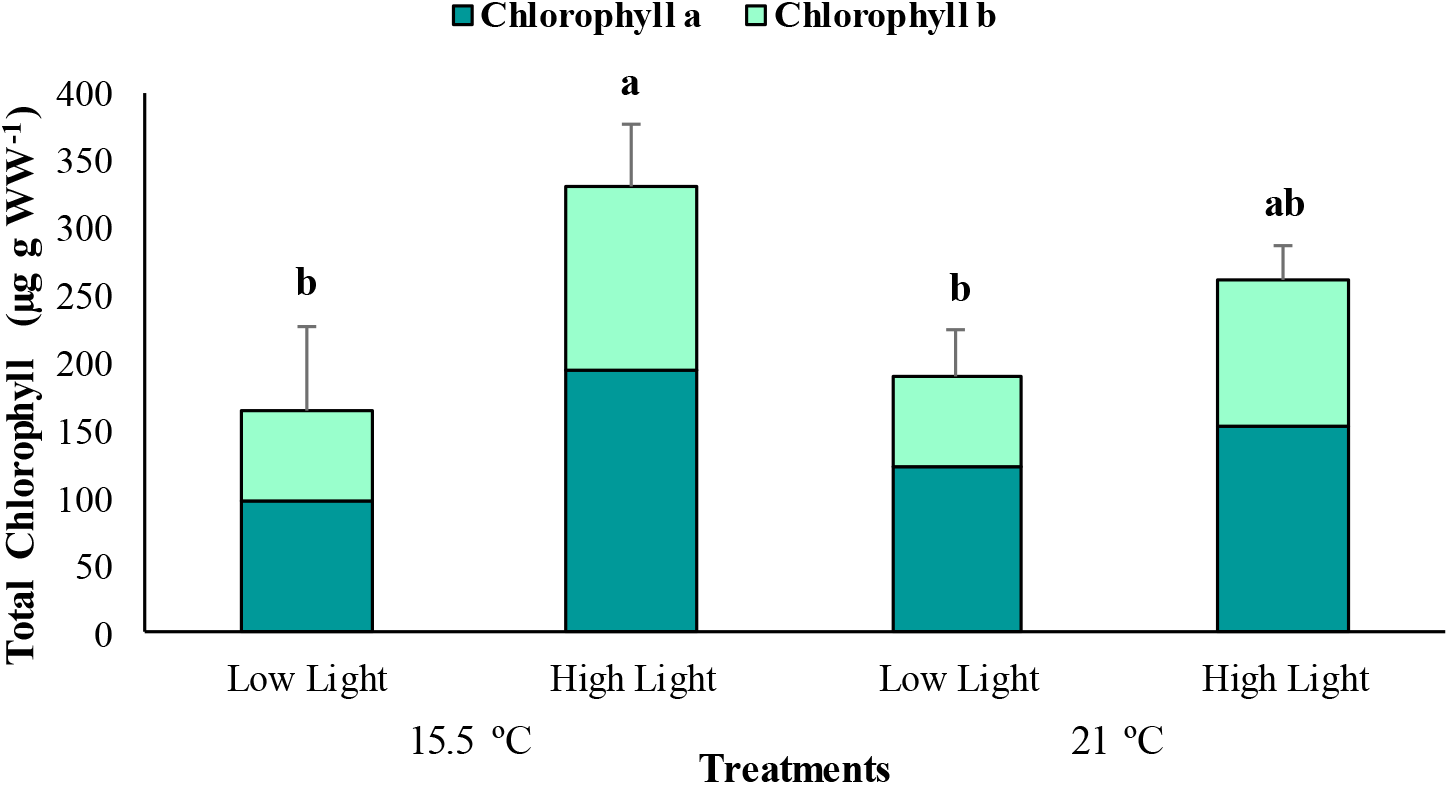
The average total chlorophyll content of *Posidonia oceanica* leaves in the different treatments (*n*=5). The error bars represent ± SD. Different letters denote significant differences (*p*<0.001) among treatments (Tukey’s post-hoc test following the analysis of deviance in Table S2, Supplementary Material).

### 3.4 Alkaline phosphatase activity

In general, APA rates differed significantly among treatments (*p*<0.05) and plant tissues (*p*<0.001) but was homogeneous between the interactions of these fixed factors (*p*>0.05; Table S3, Supplementary Material). The highest mean of APA was recorded at elevated temperature and low light conditions (5.36±4.07 μM MUF g DW^-1^ h^-1^), while the lowest corresponded to the high light treatment under equal temperature (2.80±2.26 μM MUF g DW^-1^ h^-1^). However, at the ambient temperature treatments, the values of this parameter did not deviate significantly from the rest (Fig. 5). Regarding the plant tissues, the rhizomes showed considerably lower APA rates (0.46±0.33 μM MUF g DW^-1^ h^-1^) in comparison to the leaves and roots.

**Figure 5.**
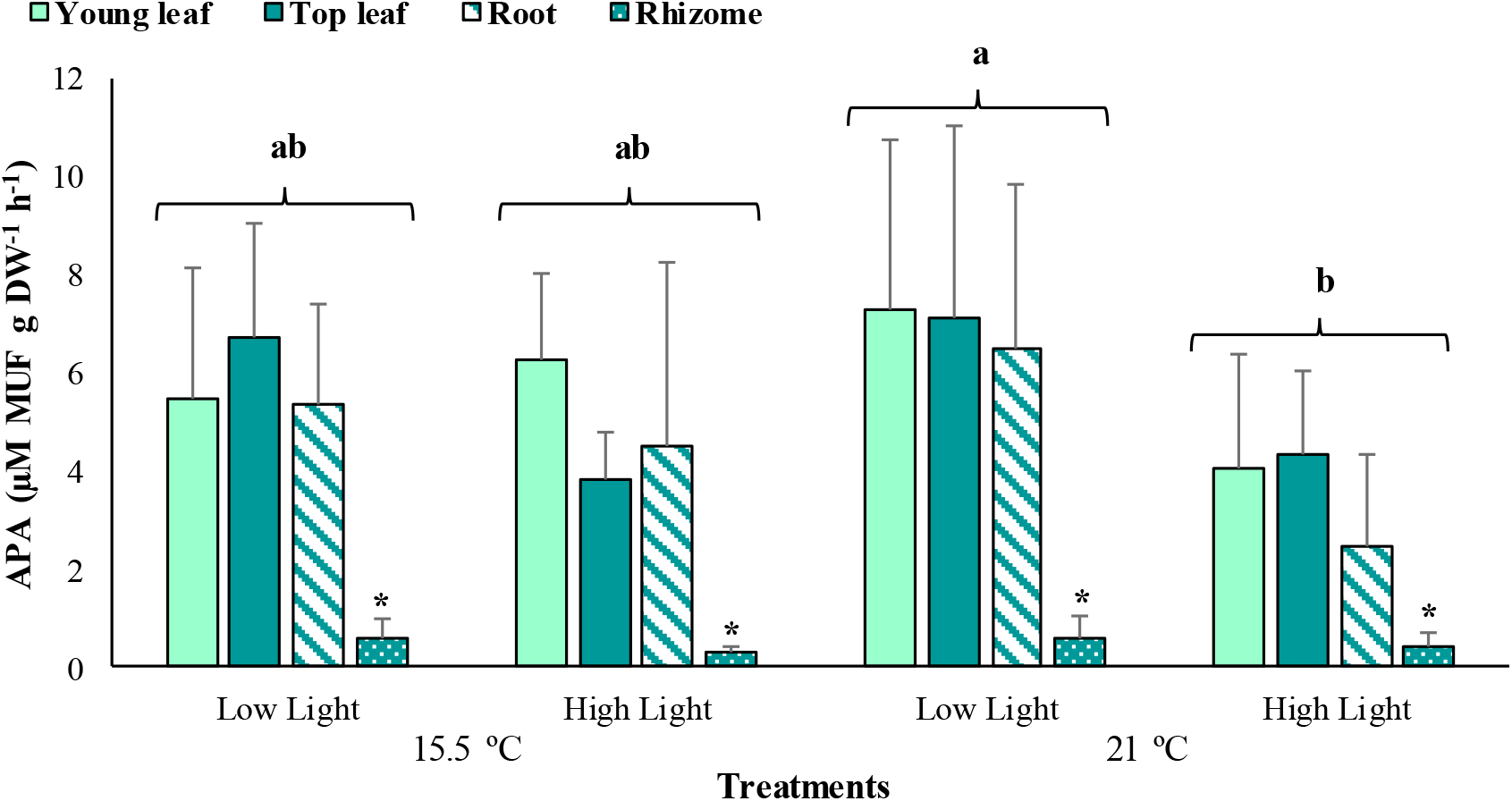
The average alkaline phosphatase activity (APA) associated with young leaves, top leaves, roots and rhizomes of *Posidonia oceanica* incubated at the different treatments (*n*=6). The error bars represent ± SD. Different letters denote significant differences (*p*<0.05) among treatments, and the asterisk (*) between plant tissues (*p*<0.001) (Tukey’s post-hoc test following the analysis of deviance in Table S3, Supplementary Material).

### 3.5 Reactive oxygen species and phenolic compounds

The reactive oxygen species (ROS) production varied significantly among treatments depending on the plant tissue (*p*<0.01; Table S4, Supplementary Material), with increased values at ambient temperature and high light conditions for the young leaves only (87.87±63.33 a.u. mg protein^-1^, Fig. 6). Additionally, it can be observed that top leaves produced greater quantities of ROS (137.54±55.70 a.u. mg protein^-1^) than the young ones (44.02±43.15 a.u. mg protein^-1^). Significant differences were also detected in the polyphenols content of *P. oceanica* between treatments depending on the plant tissue (*p*<0.05; Table S5, Supplementary Material), with the highest average amounts determined under high light conditions, but at elevated temperature and in the top leaves only (5.79±0.39 mg tyrosine mg protein^-1^, Fig. 7).

**Figure 6.**
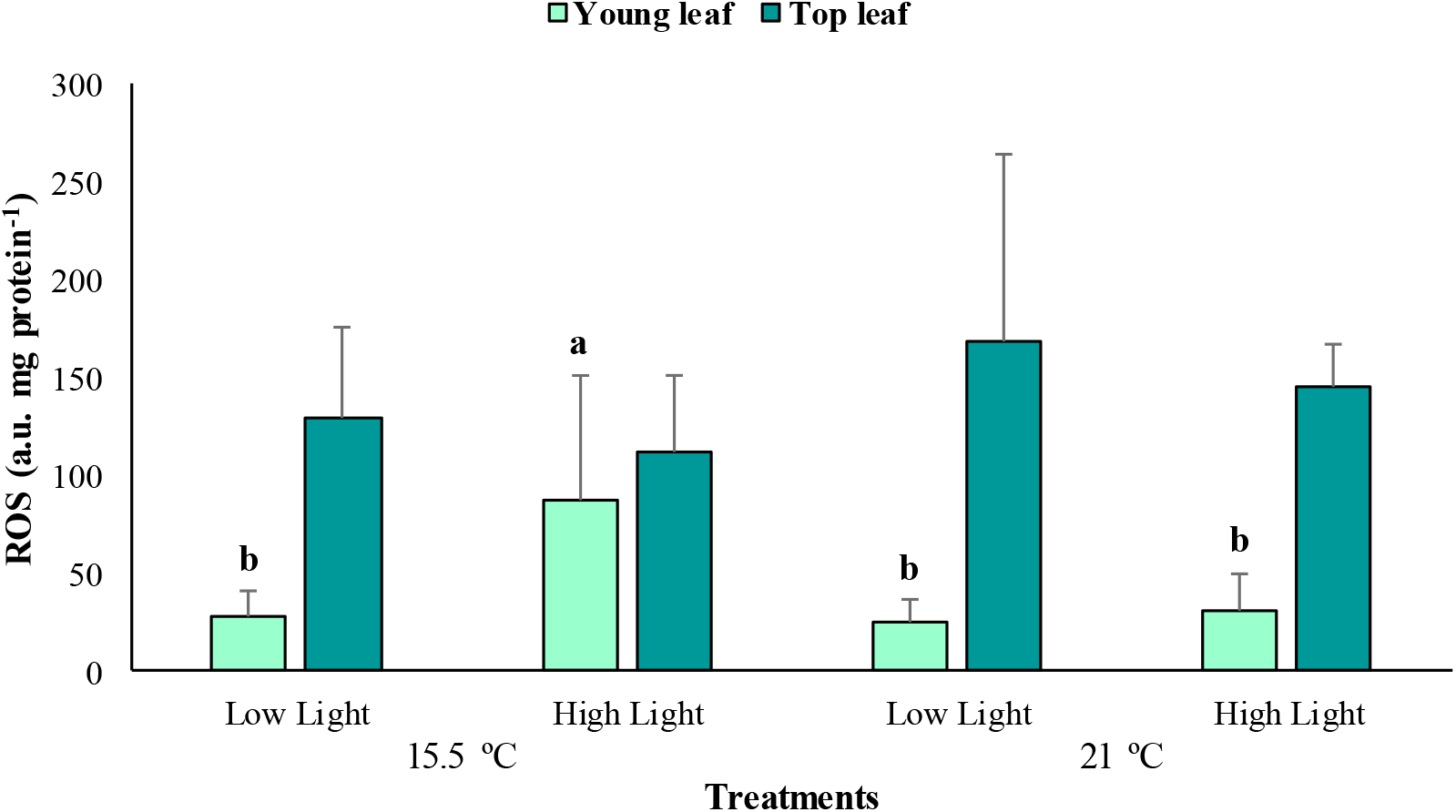
The average reactive oxygen species (ROS) production associated with young and top leaves of *Posidonia oceanica* in the different treatments (*n*=6). The error bars represent ± SD. Different letters denote significant differences (*p*<0.01) among treatments (Tukey’s post-hoc test following the analysis of deviance in Table S4, Supplementary Material).

**Figure 7.**
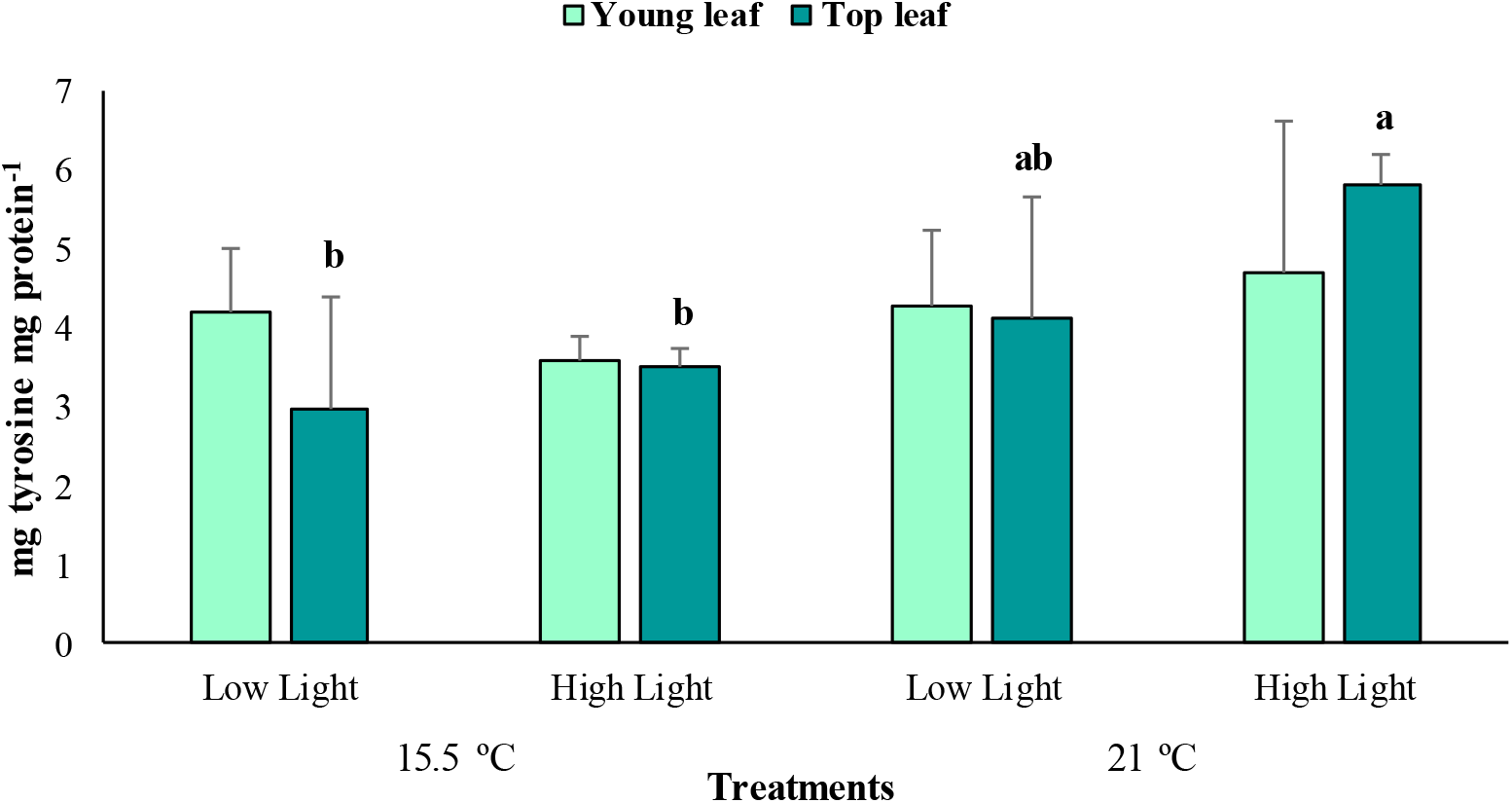
The average polyphenols content in young and top leaves of *Posidonia oceanica* in the different treatments (*n*=4). The error bars represent ± SD. Different letters denote significant differences (*p*<0.05) among treatments (Tukey’s post-hoc test following the analysis of deviance in Table S5, Supplementary Material).

### 3.6 N_2_ fixation rates and *nifH* gene expression

The estimated N_2_ fixation rates differed significantly among treatments and plant tissues (*p*<0.001; Table S6, Supplementary Material), yet the interactions between these factors did not influence the response (*p*>0.05). For the most part, the results resemble those obtained for APA, with the highest fixation rates occurring at the elevated temperature and low light treatment (0.11±0.06 nmol N_2_ g DW^-1^ h^-1^, Fig. 8), and the lowest corresponding to the treatments of ambient temperature at low light (0.02±0.03 nmol N_2_ g DW^-1^ h^-1^) and elevated temperature at high light (0.03±0.05 nmol N_2_ g DW^-1^ h^-1^). As for the plant tissues, the sterilized roots exhibited the highest N_2_ fixation rates with an average of 0.10±0.07 nmol N_2_ g DW^-1^ h^-1^, while the rhizomes demonstrated the lowest with a mean of 0.0026±0.0024 nmol N_2_ g DW^-1^ h^-1^.

**Figure 8.**
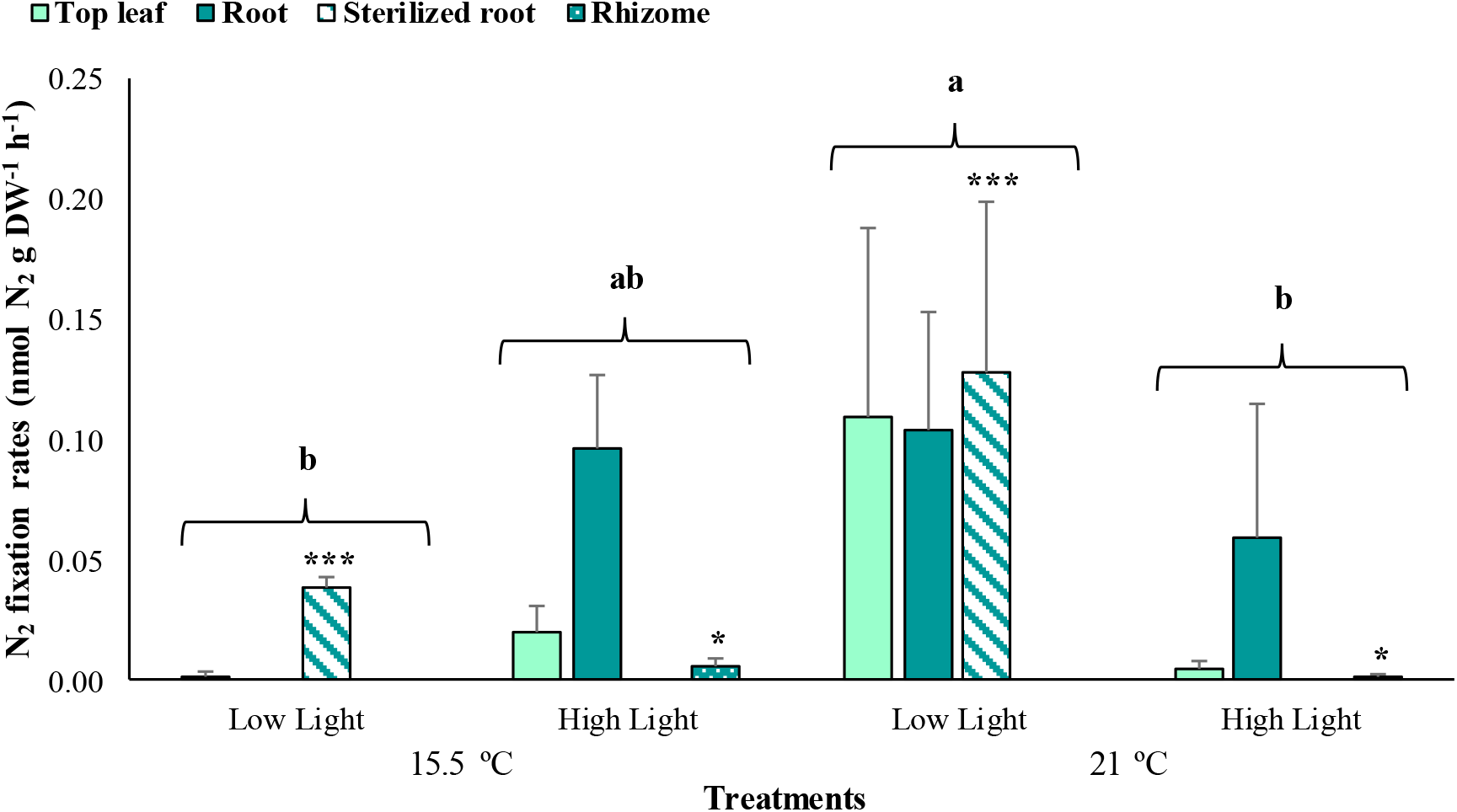
The average N_2_ fixation rates associated with young leaves, top leaves, roots and sterilized roots of *Posidonia oceanica* incubated at the different treatments (*n*=3). The error bars represent ± SD. Different letters denote significant differences among treatments, and the asterisk (*) between plant tissues (*p*<0.001) (Tukey’s post-hoc test following the analysis of deviance in Table S6, Supplementary Material).

From the groups of N_2_ fixers examined through RT-qPCR, transcripts of the *nifH* gene were only detected for the cyanobacterial phylotypes UCYN-A, -B and -C. It was determined that the total transcription of cyanobacterial groups was significantly conditioned by the type of treatment applied (*p*<0.001; Table S7, Supplementary Material). Overall, higher transcription values were obtained under elevated temperature, with UCYN-B contributing significantly with the greatest mean at low light conditions (8.46±0.32 transcripts ng total RNA^-1^, *p*<0.001, Fig. 9). For the high light treatment under equal temperature, *nifH* expression was only perceived for UCYN-B and -C. In the ambient temperature treatments, although relatively higher values were estimated at high light compared to low light, the total transcription did not differ significantly among the cyanobacterial groups (*p*>0.05). However, under low light conditions the highest *nifH* expression was attained by UCYN-C with an average of 5.65±0.36 transcripts ng total RNA^-1^, which only deviated significantly (*p*<0.01) from the mean exhibited by UCYN-A (1.22±0.82 transcripts ng total RNA^-1^). In all the treatments, the expression levels of UCYN-A were lower in comparison to the remaining groups.

**Figure 9.**
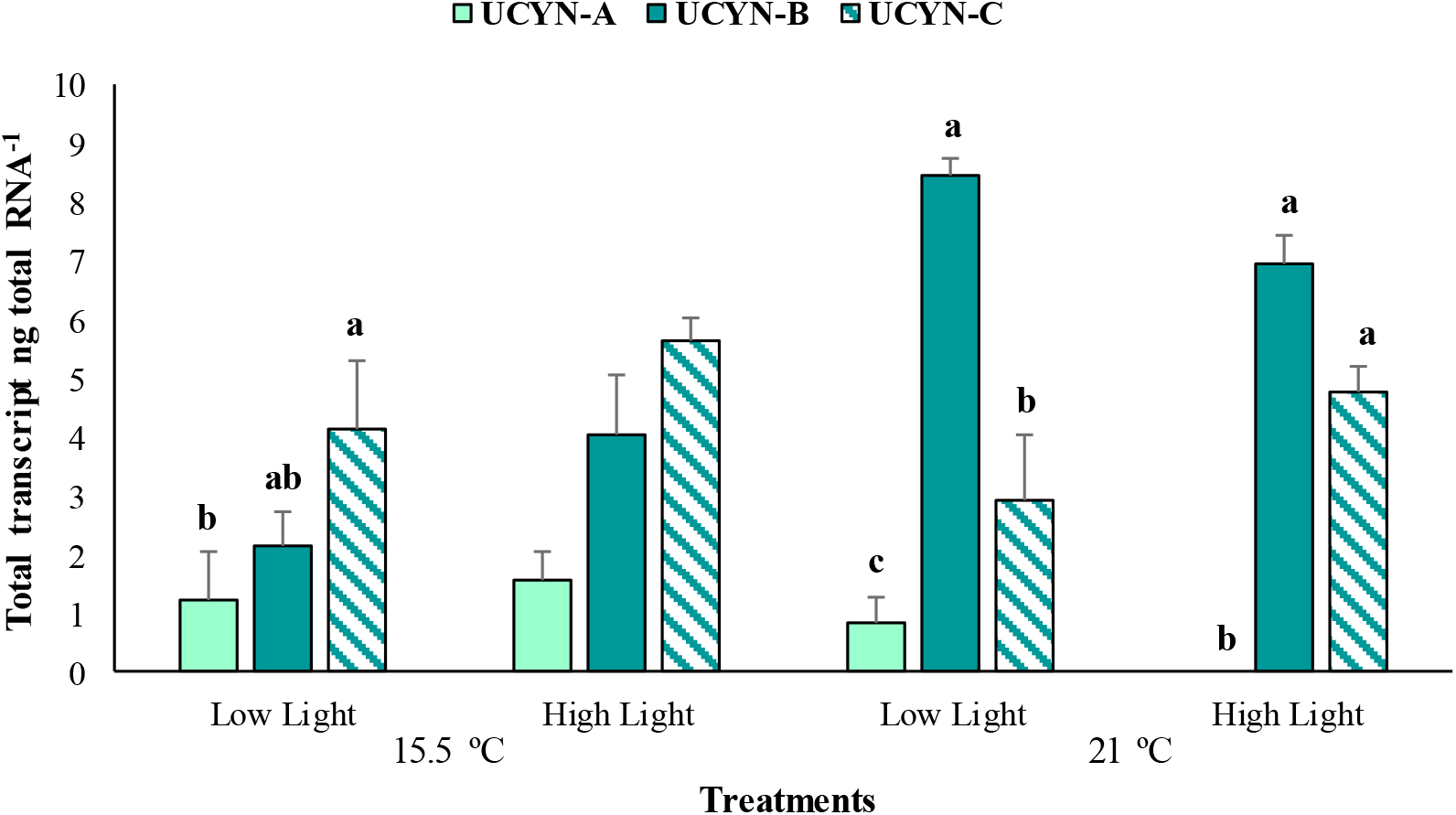
The average *nifH* gene expression of groups UCYN-A, -B and -C determined with reverse transcription quantitative polymerase chain reaction from the phyllosphere of *Posidonia oceanica* (*n*=3). The error bars represent ± SD. Different letters denote significant differences (*p*<0.001) (Tukey’s post-hoc test following the analysis of deviance in Table S7, Supplementary Material).

## 4 Discussion

The results obtained for primary productivity and chlorophyll content of *Posidonia oceanica* suggest an enhancement in these values under saturating light conditions for the plants during winter, which is in line with previous studies that emphasize light availability as the primary factor influencing the photosynthetic performance of this Mediterranean species (Pergent-Martini et al., 1994; Alcoverro et al., 1995). Higher average pH(i.e. more alkaline conditions), occurred in the aquaria under high light treatments (Fig. 2B), possibly reflecting the greater buffering effect provided by the plants through increased photosynthesis (Hendriks et al., 2013). Seagrass meadows can induce diurnal variations in the seawater carbon chemistry in relation to their productivity, generally, through the uptake of CO_2_ during photosynthesis in the day and the release of CO_2_ with respiration at night (Chou et al., 2018; Howard et al., 2018). Hence, the more metabolically intense an ecosystem is, the greater their capacity to affect the seawater pH and alkalinity (Duarte et al., 2013; Hendriks et al., 2013). In addition, greater chlorophyll production has been denoted as a photo-acclimative response of *P. oceanica* meadows thriving under high light conditions, which implies an increase in the number of reaction centers and, consequently, in the capacity of photon absorption and electron flow rate along the transport chain (Frost-Christensen and Sand-Jensen, 1992; Ruban, 2009; Dattolo et al., 2014). Higher chlorophyll content in *P. oceanica* leaves is generally related to greater photosynthetic rates (Alcoverro et al., 2001), as it was observed in this study.

During the incubations, elevated temperatures did not seem to negatively alter the photosynthetic response, nor promote higher respiration rates in *P. oceanica*, as opposed to previous findings that demonstrate the disruptive effect of this factor on the productivity of this seagrass species (Collier and Waycott, 2014; Marín-Guirao et al., 2018). This could be partially attributed to the fact that, although the elevated temperature treatment applied in the aquaria was 1°C above the optimum conditions recorded for this seagrass (17-20°C), it stays within its temperature comfort range (13-24°C) (Boudouresque and Meinesz, 1982). However, significantly higher leaf net photosynthetic rates were still exhibited by plants subjected to 15.5 °C and high light treatment. On the other hand, in Agawin et al. (2021), positive responses to higher temperature were observed in the photosynthetic activity of *P. oceanica*, while leaf respiration rates did not increase, suggesting that ocean warming scenarios may not necessarily have adverse effects on the carbon balance of the plants in winter. This agrees with the higher NPP rates estimated from the phyllosphere of *P. oceanica* at elevated temperature and high light conditions during the experiment.

The results of this study showed a positive response in the productivity of *P. oceanica* to light availability in winter. This suggests that, anthropogenic activities that cause prolonged periods of reduced surface irradiance will possibly have more destructive impacts in these ecosystems during winter, rather than the prospected sea warming in the Mediterranean. Similarly, Hendriks et al. (2017) reported that low light availability had a negative effect on the photosynthetic performance of *P. oceanica* under short-term experimental conditions for summer, while temperature negatively affected the plants growth. Champenois and Borges (2018) also highlighted the strong association between the GPP of *P. oceanica* and the interannual variations of light availability over a decade in Bay of Revellata, France, and a positive correlation to temperature, given that the temperatures recorded stayed within the comfort range of the seagrass species. Nevertheless, Serrano et al. (2011) demonstrated through *in situ* shading experiments in Portlligat Bay, Spain, that shallow *P. oceanica* meadows are more vulnerable to severe light limitation during spring-summer compared with autumn-winter, since it coincides with the plants’ favorable growth season when they accumulate reserves for overwintering.

The biochemical responses of *P. oceanica* showed that young leaves subjected to the ambient temperature and high light treatment exhibited significantly higher ROS production, while top leaves demonstrated greater polyphenols content at elevated temperature. These observations coincide with studies that report how factors such as high irradiance and temperature may promote the production of ROS in photosynthetic organisms, that in excessive quantities leads to oxidative stress (Choo et al., 2004; Adams et al., 2006; Costa et al., 2015). In this study, the young leaves of *P. oceanica*, that were still in the process of development, might have been more vulnerable to the saturating light conditions, which possibly induced the elevated ROS concentrations detected. However, it should also be noted that high primary production rates unavoidably prompt the production of ROS (Hajiboland, 2014), since these free oxygen radicals are liberated when the photolysis of water molecules by the photosystem II (PSII) occurs during photosynthesis (Lesser, 2006). This corresponds with the significantly higher NPP rates reported for the young leaves at the ambient temperature and high light treatment, hence, the increased amounts of ROS perceived in these tissues could also be related to their high photosynthetic activity. For the top leaves, the higher phenolic compounds measured could be a response of the plants to the elevated temperatures they were exposed to, considering they were not optimum for their functioning and seagrasses, especially for *P. oceanica* which are sensitive to the quality of environmental conditions (Orth et al., 2006). Phenolic compounds demonstrate several biological functions that include antioxidant activity, and plants generally activate antioxidant mechanisms to detoxify the ROS generated and avoid oxidative stress (Cheynier et al., 2013; Costa et al., 2015). The higher mean values of ROS calculated for top leaves under elevated temperature compared to ambient treatments may support this notion, although the differences were not significant (*p*<0.05).

The phosphatase activity in *P. oceanica* achieved its maximum values at elevated temperature and low light conditions, which is consistent with the patterns described by Invers et al. (1995), who demonstrated that increasing temperatures can positively affect APA rates in this seagrass until a certain threshold (24 °C). The high activity of this enzyme under elevated temperature and low light also matches the significantly enhanced N_2_ fixation rates exhibited by this treatment, considering that the energy, in the form of adenosine triphosphate (ATP), to fuel N_2_ fixation is dependent on the presence of inorganic phosphorus, therefore, the demand for this nutrient is theoretically induced when the cells are fixing N_2_ (Romano et al., 2017; Fernández-Juárez et al., 2019). In contrast, the APA values were on average the lowest under equal temperature and saturating light, regardless of the more favorable pH values for the phosphatases in this treatment, as they were higher compared to the low light treatments. This result could be attributed to the fact that at the final stage of the experiment the mean phosphate concentration for this treatment was several orders of magnitude higher than the rest (Table 1), given that APA in seagrasses decreases under elevated phosphate content and, vice versa, increases with phosphorus limitation (Invers et al., 1995; Martínez-Crego et al., 2006; Agawin et al., 2021). As for the significantly lower APA rates demonstrated by the rhizomes, this may suggest that the metabolic activity occurring in this part of the plants is relatively low compared to the others, consequently, its inorganic phosphorus demand is low as well. Generally, the phosphatase activity tends to be greater in the leaves, partially due to the contribution of the epiphytes (Invers et al., 1995).

The N_2_ fixation rates estimated in the present study are relatively similar to those previously obtained by other works in *P. oceanica* meadows during winter (Agawin et al., 2017, 2019). Diazotrophic activity being significantly higher at elevated temperature and low light conditions could be attributed to the strong temperature dependency of the nitrogenase enzyme (Brauer et al., 2013; Agawin et al., 2017; Garcias-Bonet et al., 2019). Further, the low nitrate concentrations exhibited in this treatment at the final stage of the incubations (Table 1) may suggest the existence of dissolved inorganic nitrogen limitation, which possibly induce N_2_ fixing conditions in these aquaria. Nonetheless, N_2_ fixation was considerably lower under elevated temperature and saturating light conditions and this could be related to the significantly high GPP values measured for the leaves and phyllosphere of *P. oceanica* in these tanks. It has been widely documented that N_2_ fixation is an oxygen sensitive process, since molecular oxygen (O_2_) is capable of inactivating the nitrogenase and causing irreversible damage to the protein structure, as well as inhibiting the synthesis of the enzyme in many diazotrophs (Berman-Frank et al., 2003; Schoffman et al., 2016). Thus, increased O_2_ evolution with increased photosynthesis may affect negatively the N_2_ fixation activities associated with *P. oceanica* in treatments with increased GPP. Diazotrophic activity showed variability among plant parts, with the roots exhibiting higher average values, particularly the sterilized ones containing the root endophytes, in comparison to the leaves. These findings are consistent with the results of Lehnen et al. (2016) for *P. oceanica* and Hamisi et al. (2009) in tropical seagrass species. According to the latter authors, higher activities in the rhizosphere could be associated with a high occurrence of heterotrophic diazotrophs aside from the autotrophic bacteria in the phyllosphere. This pattern is also in agreement with the reported by Agawin et al. (2019), who determined the presence of seasonality in the N_2_ fixation process related to *P. oceanica* meadows along the Mallorcan coast, with generally higher activities associated with the roots during winter.

The RT-qPCR analyses revealed the presence of the three groups of unicellular diazotrophic cyanobacteria in the phyllosphere of *P. oceanica*, with UCYN-B and -C displaying notably higher transcription levels of the *nifH* gene in comparison to UCYN-A. Past molecular analyses carried out by Agawin et al. (2017) indicated the presence of members of the UCYN-B, such as *Crocosphaera*, and UCYN-C, like *Cyanothece*, in addition to other genera of the phyla Cyanobacteria, Proteobacteria, Firmicutes, Bacteroidetes, and Archaea in the phyllosphere of *P. oceanica*. The UCYN-A (*Candidatus Atelocyanobacterium thalassa*) is regarded as one of the most abundant and widespread N_2_ fixing groups in the ocean, proved to live attached or in symbiosis with larger single-celled prymnesiophytes, given that they lack important biosynthetic pathways genes, including oxygenic photosynthesis and carbon fixation (Zehr et al., 2008; Thompson et al., 2012). Considering its association with larger algae and the frequent symbioses between cyanobacteria and multicellular plants evidenced in terrestrial habitats (Thompson et al., 2012), the presence of this N_2_ fixing group in *P. oceanica* is also plausible. The diazotrophic communities identified showcased varied responses to the treatments, with UCYN-B demonstrating the maximum average of *nifH* expression overall at elevated temperature and limited light, while UCYN-C had significantly higher values compared to the rest under ambient temperature and low light conditions, although the highest means for this latter group alone were under both high light treatments. The UCYN-B group having enhanced activities at elevated temperatures is consistent, based on the literature (Brauer et al., 2013; Agawin et al., 2017), with the general positive correlation recorded between temperature and N_2_ fixation. Moreover, past studies have evidenced the thermophilic habit of *Crocosphaera* (UCYN-B), with warmer sea surface temperatures (26-29 °C) primarily determining its distribution across the oceans (Church et al., 2008; Moisander et al., 2010). The results obtained for UCYN-C might be due to the different light and temperature requirements its representatives possess, consequently, the species have optimum N_2_ fixation rates under distinct conditions. On the other hand, although it has been denoted that UCYN-A exhibits a broad temperature range, more associated with cooler waters and a lower temperature optimum than the remaining cyanobacterial groups (Moisander et al., 2010; Cabello et al., 2020), the results obtained from the experiment do not reflect a clear pattern. Taking into consideration how the different diazotrophic species are adapted to grow and function under differing conditions, further investigation is required to achieve better understanding of their potential response to the interaction between climate change factors and other stressors.

## 5 Author Contributions

MGG-M and NA designed the experiments. MGG-M, VF-J and JCR-C conducted all experiments. All authors led the writing of the manuscript, reviewed, and supervised by the head of the laboratory, NA.

## 6 Funding

This work was supported by funding to NA through the Ministerio de Economía, Industria y Competitividad-Agencia Estatal de Investigación, and the European Regional Development Funds project (CTM2016-75457-P).

## 7 Acknowledgments

We acknowledge help of the Scientific Technical Service of the University of the Balearic Islands in the gas chromatography (Maria Trinidad Garcia Barceló), spectrofluorimetric (Biel Martorell Crespí) and flow injection analyses (Josep Agustí Pablo Cànaves). We also thank the support of Regino Martinez (Spanish Institute of Oceanography) in the ammonium analysis.

## Notes

### Competing Interest Statement

The authors have declared no competing interest.

