## Supplementary Material for "Response of *Posidonia oceanica* (L.) Delile and its associated N_2_ fixers to different combinations of temperature and light levels"

**Supplementary Table 1.** Univariate analysis of variance (ANOVA) to test the effect of different combinations of light levels and temperature on the gross primary production (GPP), net primary production (NPP) and respiration rates of *Posidonia oceanica* leaves; and analysis of deviance for the phyllosphere data set.

| **Response** | **Source** | **Df** | **Sum Sq** | **Mean Sq** | **F value** | **Pr(>F)** |
|---|---|---|---|---|---|---|
| GPP | Treatment | 3 | 33.70 | 11.23 | 5.71 | 0.02185* |
| | Residuals | 8 | 15.75 | 1.97 | | |
| NPP | Treatment | 3 | 18.98 | 6.33 | 6.17 | 0.0178* |
| | Residuals | 8 | 8.21 | 1.03 | | |
| Respiration | Treatment | 3 | 6.21 | 2.07 | 0.46 | 0.7185 |
| | Residuals | 8 | 36.10 | 4.51 | | |
| logGPP (phyllosphere) | | | | | **Chisq** | **Pr(>Chisq)** |
| | Treatment | 3 | | | 14.32 | 0.002498** |

*Significant at p<0.05; **very significant at p<0.01.

**Supplementary Table 2.** Analysis of deviance to test the effect of different combinations of light levels and temperature on the total chlorophyll, along with chlorophyll *a* and *b*, of *Posidonia oceanica* leaves.

| **Response** | **Source** | **Chisq** | **Df** | **Pr(>Chisq)** |
|---|---|---|---|---|
| Total Chl | Treatment | 20.41 | 3 | 0.0001395*** |
| Chl *a* | | 18.91 | 3 | 0.0002849*** |
| Chl *b* | | 18.82 | 3 | 0.0002979*** |

***Highly significant at *p*<0.001.

**Supplementary Table 3.** Analysis of deviance to test the effect of different combinations of light levels and temperature on alkaline phosphatase activities associated with the young leaves, top leaves, roots and rhizomes of *Posidonia oceanica*.

| **Source** | **Chisq** | **Df** | **Pr(>Chisq)** |
|---|---|---|---|
| Treatment | 9.76 | 3 | 0.02077* |
| Plant Tissue | 463.55 | 3 | < 0.0001*** |
| Treatment × Plant Tissue | 10.98 | 9 | 0.27713 |

*Significant at *p*<0.05; ***highly significant at *p*<0.001.

**Supplementary Table 4.** Analysis of deviance to test the effect of different combinations of light levels and temperature on reactive oxygen species production associated with young and top leaves of *Posidonia oceanica*.

| Source | Chisq | Df | Pr(>Chisq) |
|---|---|---|---|
| Treatment | 4.95 | 3 | 0.17534 |
| Plant Tissue | 76.99 | 1 | < 0.0001*** |
| Treatment × Plant Tissue | 15.38 | 3 | 0.001516** |

**Very significant at $p<0.01$; ***highly significant at $p<0.001$.

**Supplementary Table 5.** Analysis of deviance to test the effect of different combinations of light levels and temperature on the polyphenols content of young and top leaves of *Posidonia oceanica*.

| Source | Chisq | Df | Pr(>Chisq) |
|---|---|---|---|
| Treatment | 6.54 | 3 | 0.08827 |
| Plant Tissue | 0.00 | 1 | 0.98606 |
| Treatment × Plant Tissue | 9.24 | 3 | 0.02632* |

*Significant at $p<0.05$.

**Supplementary Table 6.** Analysis of deviance to test the effect of different combinations of light levels and temperature on the $N_2$ fixation rates associated with the young leaves, top leaves, roots and sterilized roots of *Posidonia oceanica*.

| Response | Chisq | Df | Pr(>Chisq) |
|---|---|---|---|
| Treatment | 16.77 | 3 | 0.0007885*** |
| Plant Tissue | 19.34 | 3 | 0.0002321*** |
| Treatment × Plant Tissue | 3.93 | 4 | 0.4150302 |

***Highly significant at $p<0.001$.

**Supplementary Table 7.** Analysis of deviance to test the effect of different combinations of light levels and temperature on the *nifH* gene expression of groups UCYN-A, -B and -C associated with the phyllosphere of *Posidonia oceanica*.

| Source | Chisq | Df | Pr(>Chisq) |
|---|---|---|---|
| Treatment | 12.94 | 3 | 0.00478** |
| Group | 216.97 | 2 | < 0.0001*** |
| Treatment × Group | 91.40 | 6 | < 0.0001*** |

***Highly significant at $p<0.001$.
